# Geometric causes of species rarity

**DOI:** 10.64898/2026.09.02.748835

**Authors:** Anna Toszogyova, David Storch

## Abstract

Understanding the limits of species distributions is a central objective of biogeography and macroecology and has become increasingly important as climate change drives rapid shifts in geographic ranges. Species range sizes follow a highly skewed frequency distribution, with most species occupying ranges orders of magnitude smaller than those of the most widespread species. Range sizes also exhibit pronounced geographic patterns, with small-ranged species concentrated near continental margins and other geographic boundaries. No universally accepted explanation has been proposed for these patterns. Here we present a simple geometric model showing that species range size patterns emerge from the random placement of dispersal barriers within continental domains. The model predicts both the observed frequency distribution and the spatial distribution of range sizes across amphibians, birds, and mammals. It therefore provides a first-order explanation for global patterns of species rarity and can be refined by incorporating elevational barriers and spatial variation in species richness. Our findings suggest that species range size is constrained by the geometry of dispersal barriers and the geographic domain, with proximity to domain boundaries acting as a primary determinant of species rarity. These results have important implications for understanding species’ evolutionary potential and vulnerability to extinction.

---

Species geographic ranges represent a natural unit of macroecological analyses, yet causes underlying the patterns in range sizes are surprisingly poorly understood (1–3) and the question why so many species have very restricted geographic distribution remains open. Similarly to the species abundance distribution (SAD), the range size frequency distribution is typically close to lognormal, most ranges being orders of magnitude smaller than a few largest ranges (4–7), but in contrast to the SAD, attempts to explain the range size frequency distribution have been surprisingly rare (8, 9). Also, even though patterns comprising global distribution of range sizes have been reported, general trends are ambiguous. Specifically, although Rapoport’s rule (10) states that range size should decrease towards the tropic, empirical patterns appear to be more complex, with a tendency of range size to decrease from high latitudes of the northern hemisphere towards southernmost parts of the southern hemisphere (11–13), range size being strongly affected by the size of respective geographic domain (11). Smaller ranges are typically found in tropical and subtropical mountain ranges and along coasts, peninsulas, and insular regions (11, 13). Understanding the processes underlying these patterns is crucial for interpreting biodiversity dynamics, anticipating species’ responses to environmental changes, and identifying regions most at risk of biodiversity loss. This is particularly important because extinction risk depends on range size and the spatial position of species ranges, with species being lost more rapidly when habitat loss proceeds from region boundaries inward (14), highlighting the potential importance of geometry and domain edges in shaping both species rarity and vulnerability.

## Geometric model

Here we propose a first-order model predicting elevated species rarity along continental margins (see also ref. 13) and an overall prevalence of small ranges. The model is based on an assumption that dispersal barriers and their distribution across continents represent key determinants of range sizes. According to the model, there are two principally different kinds of barriers: hard outer boundaries of land masses (i.e. coastlines) that are shared by all species, and inner barriers that are species-specific and may correspond to various dispersal limits including mountain ridges, edges of suitable habitats or presence of competitors. To illustrate the idea using a simplified square-shaped land mass, let’s assume that within this domain, the inner barriers are vertical and horizontal lines whose position is random (i.e. their intersections with the horizontal and vertical axes are drawn from a uniform distribution; Fig. 1). A species originates at any point within the domain and spreads until it reaches the nearest barriers that define its range. Species which originate close to the domain edge are constrained in that direction and can only spread in other directions. However, since the position of the inner barriers is independent of the outer boundaries, there is a considerable chance that the species will encounter an inner barrier shortly thereafter, and will remain trapped between the barriers, resulting in a small range size. In contrast, species which originate near the center of the domain have a higher probability of being able to spread further in at least some directions (Fig. 1A). As a result, the model predicts that average range size decreases toward the edges of the domain (Fig. 1 B and C). The frequency distribution of range sizes generated by the model is characterized by prevalence of small ranges and is nearly lognormal (Fig. 1D), as most ranges are constrained by the barriers that lie relatively close to each other, and there is only a low probability that the random position of the barriers creates a large space between them. The exact properties of both patterns depend on only one free parameter, namely the density of inner barriers.

**Fig. 1:**
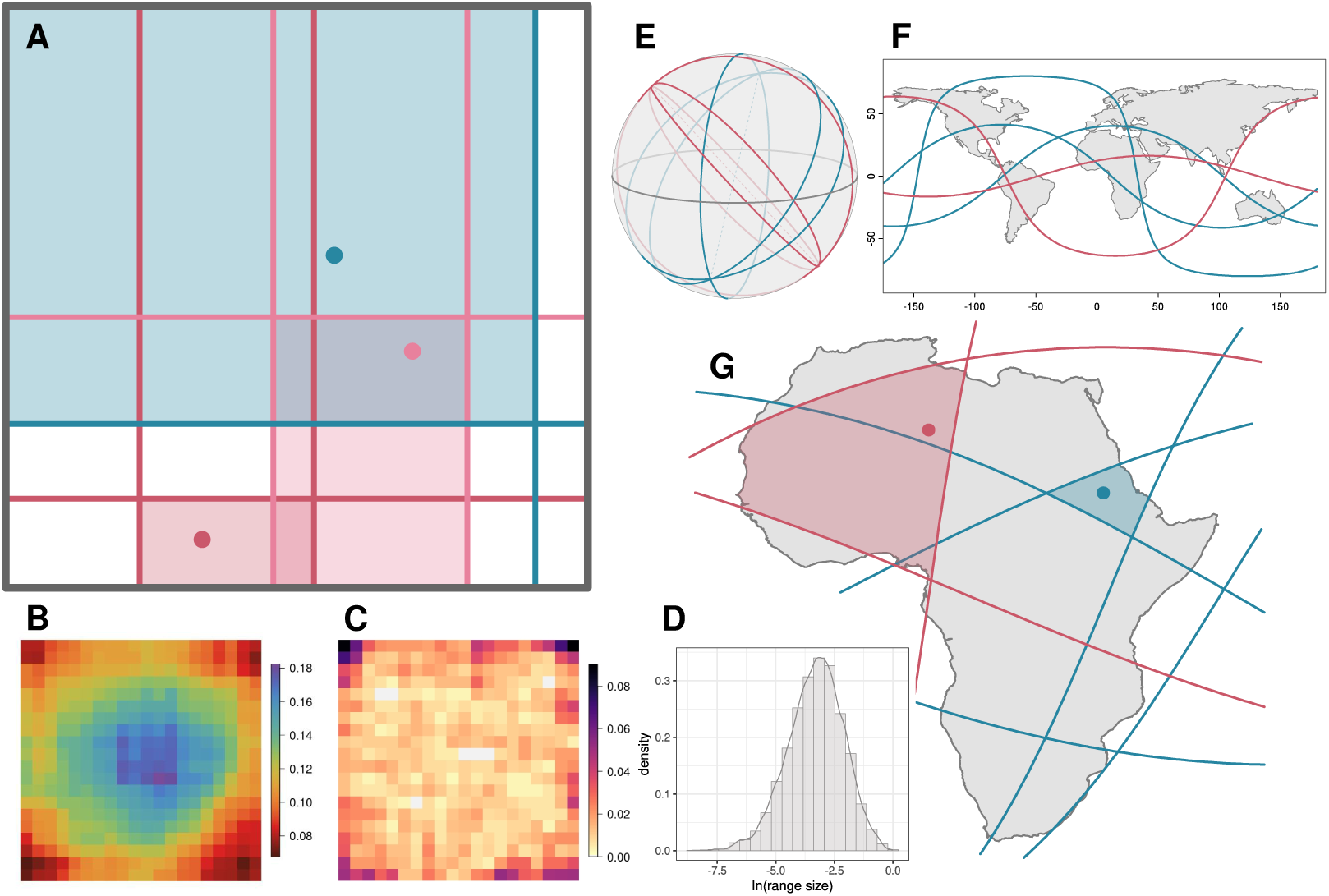
Geometric model of species ranges. (A) Schematic representation of the model based on random placement of dispersal barriers. The edges of the square domain define hard outer boundaries shared by all species. Inner barriers (randomly positioned vertical and horizontal lines, specific for each species) limit species spread from randomly located points of origin until the nearest barriers are reached. Species ranges originating near domain edges are constrained by the domain edge and are likely to be further limited by species-specific inner barriers, resulting in smaller ranges. This leads to particular spatial patterns of median range size (B) and of the proportion of small-ranged species (first 15th percentile) (C). Patterns in (B) and (C) are shown on a 20 × 20 grid. (D) Frequency distribution of range sizes generated by the model (log scale). Results for (B to D) are based on simulations of 5,000 species with inner-barrier density Pois(*D* = 7). (E) The globe with randomly placed inner barriers represented by great circles. (F) Continental domains and respective inner barriers in WGS84 projection. (G) An example for Africa, where the species originate at randomly located points and expand until reaching the nearest barriers. Two different colors refer to two species with species-specific inner barriers.

## Spherical surface implementation

Using spherical geometry comprising the Earth’s surface, we apply this conceptual framework to real geographic domains (i.e. continents), modeling the inner barriers within continents (Eurasia, Africa, North America, South America, and Australia) as randomly placed great circles (the circles whose planes pass through the sphere’s center; Fig. 1E–G). The model generates conspicuous geographic patterns of range size, consisting in the increasing prevalence of small ranges towards continental edges (Fig. 2A–F), in accordance with the observed patterns (Fig. 2G–I and *SI Appendix*, Fig. S1). Model outputs depend on the density of inner barriers (Fig. 2A–F), typically with the best performance at inner-barrier densities of 15–25. The fit varied among continents and taxa, explaining about 60% of spatial variation in median range size in Australia for all taxa, while in South America the model captured 24%, 33%, and 35% of variation for birds, mammals, and amphibians, respectively. In North America, the best fit was obtained for birds (40%), and in Africa for amphibians (29%). The lowest portion of variation was captured by the model in Eurasia (Table 1).

**Fig. 2:**
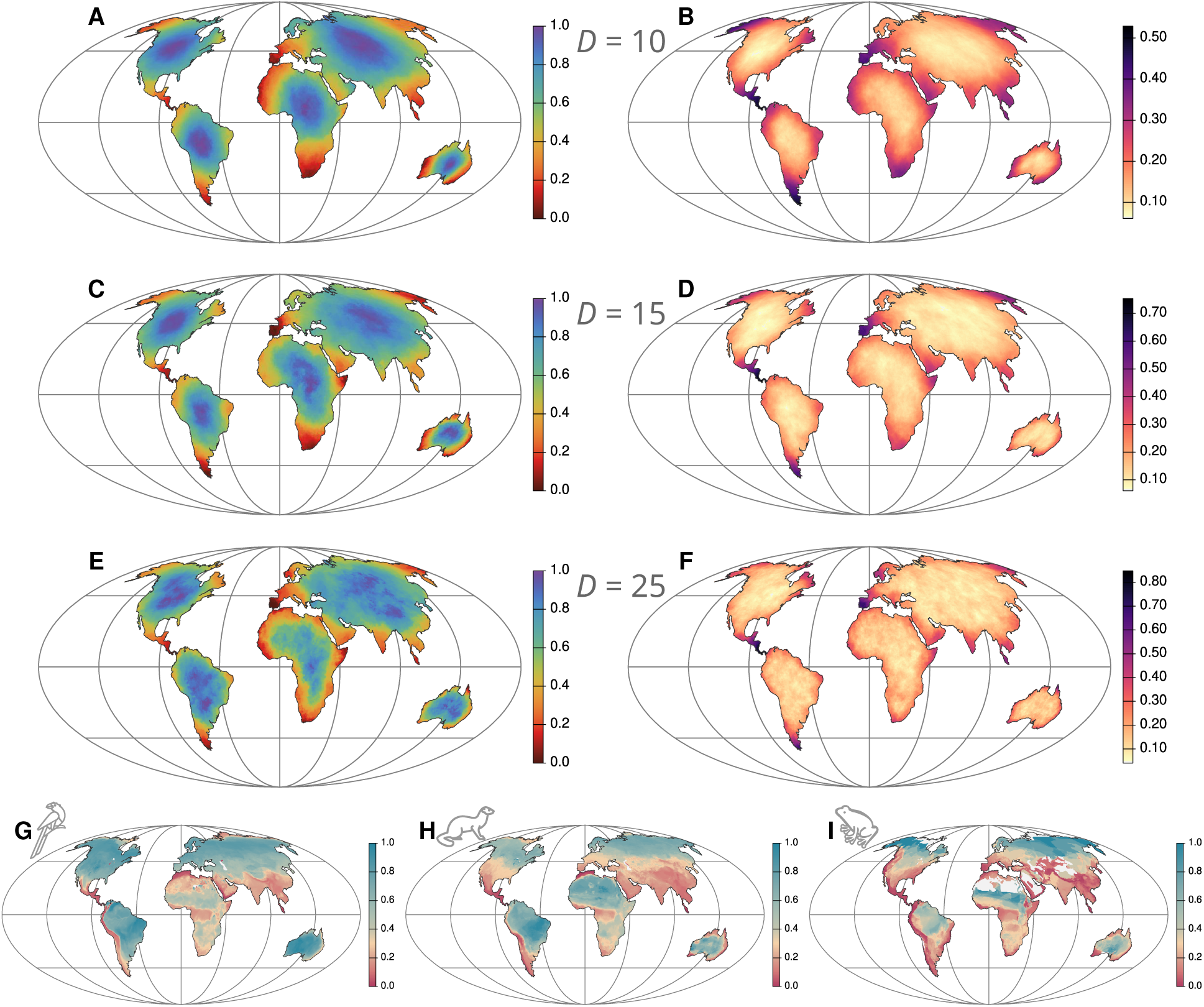
Modeled and observed geographic patterns of range size. (A, C, E) Geographic patterns of modeled median range size under varying barrier densities *D*, scaled separately within each continent (red indicates smaller ranges, blue larger ranges). (B, D, F) Geographic patterns of the proportion of small-ranged species (below the continental median). The results are based on simulations of 5,000 species and rasterized to 30-km grid cells in the Mollweide projection. Inner barriers are placed randomly with densities Pois(*D* = 10) (A and B), Pois(*D* = 15) (C and D), and Pois(*D* = 25) (E and F). (G, H, I) Geographic patterns of median range size for birds (G), mammals (H), and amphibians (I). The maps are displayed in the Mollweide projection and rasterized at 30-km resolution.

**Table 1:**
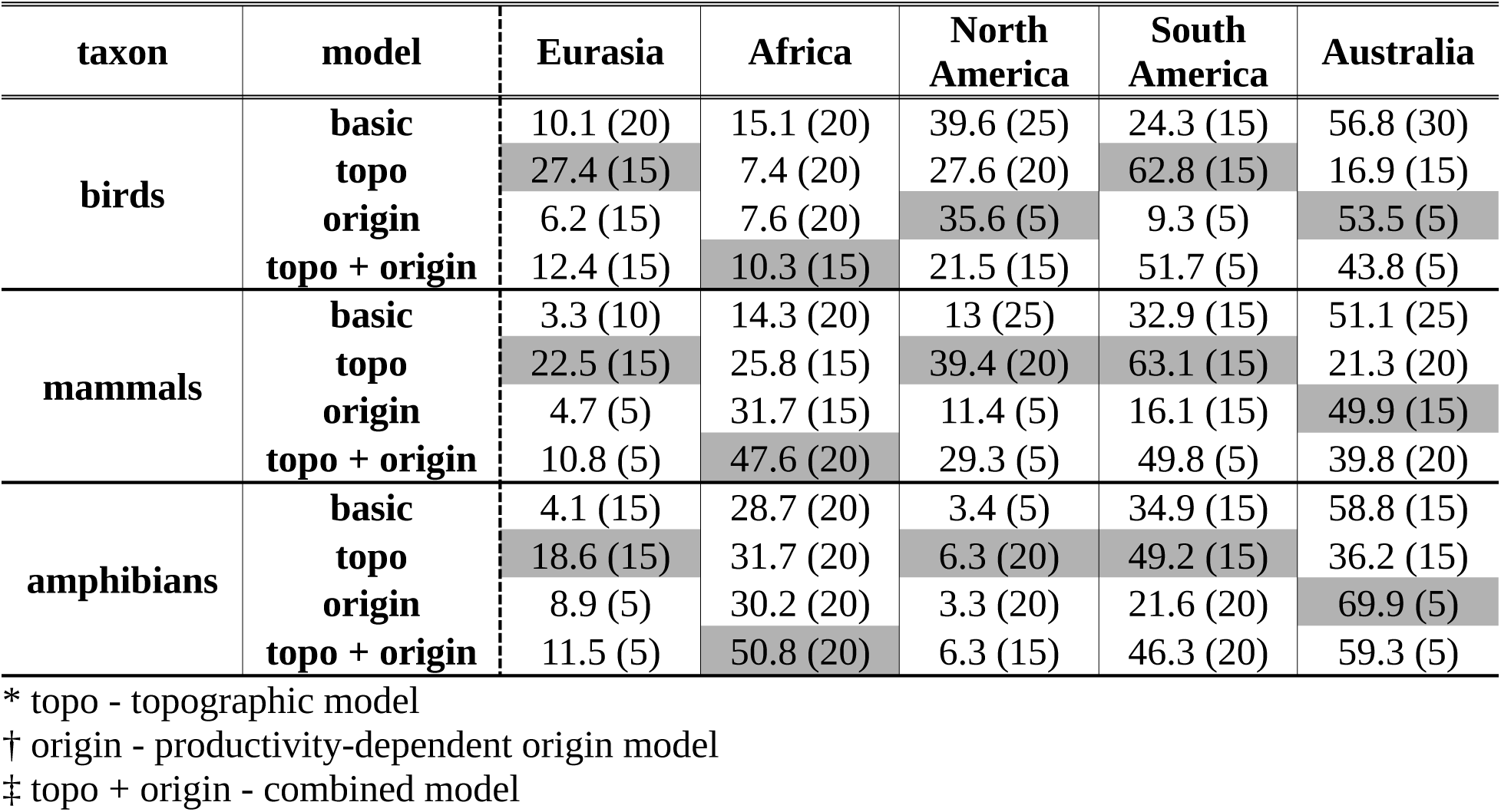
Variance in median range size explained by the geometric model. Adjusted coefficients of determination (R2, %) from linear models relating observed median range size across grid cells (dependent variable) to model-predicted median range size (explanatory variable) for each taxon and continent. The models with best-performing densities of inner barriers for each combination have been selected; the density of inner barriers for these models is shown in parentheses. The best-performing values within the advanced models (those that include elevation and/or productivity-dependent species origination probability) are highlighted in grey.

The frequency distribution of range sizes within individual land masses generated by the model was close to lognormal, although a bit left-skewed on the logarithmic scale, similarly to the observed distributions (Fig. 3). The model slightly overestimated mean range sizes, especially at lower barrier densities, and failed to reproduce the smallest ranges (*SI Appendix*, Table S1). The observed frequency distributions reveal higher variability among taxa and land masses than model outputs, amphibians having generally smaller ranges and considerably flatter frequency distributions than other taxa and model predictions. This indicates that although the model captures general properties of the patterns, there are additional constraints on species ranges which may be linked to biological traits or evolutionary history of respective taxa.

**Fig. 3:**
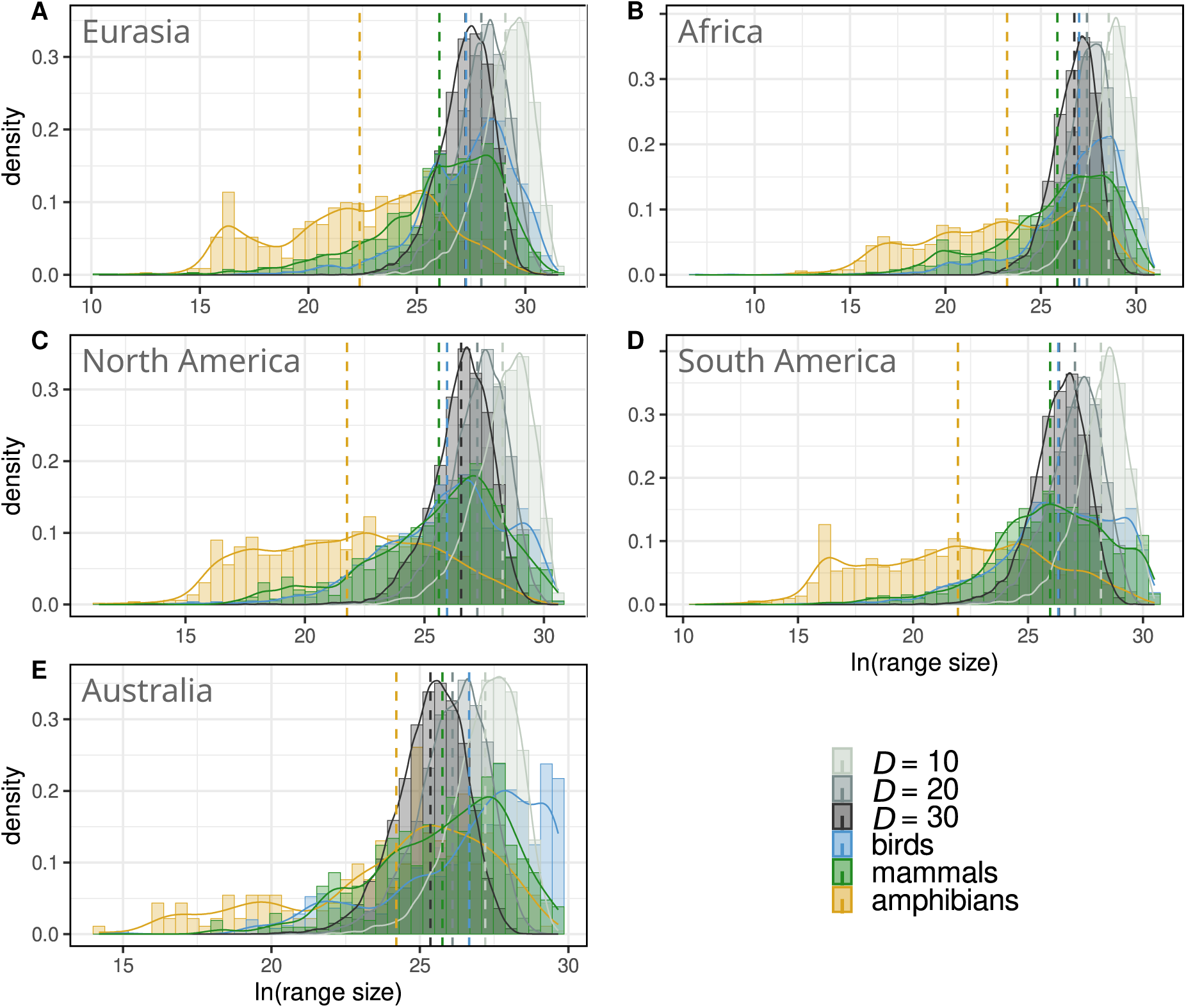
Frequency distributions of range sizes predicted by the basic geometric model and compared with the observed distribution for three taxa. Frequency distributions of modeled (grey) and observed range sizes for birds (blue), mammals (green), and amphibians (yellow) in Eurasia (A), Africa (B), North America (C), South America (D), and Australia (E), shown on a logarithmic scale (m^2^). Modeled distributions are presented for three inner-barrier densities: Pois(*D* = 10) (light grey), Pois(*D* = 20) (grey), and Pois(*D* = 30) (dark grey). Both observed and modeled distributions are approximately lognormal and left-skewed on the log scale, with modeled means generally exceeding observed values. Modeled distributions also exhibit higher kurtosis. Increasing barrier density reduces skewness and lowers mean range size. Solid lines represent kernel densities, and dashed lines indicate mean values.

## Adding realism to the models

### Topographic model

To increase the realism of the model, we incorporated topography by an addition of barriers driven by an elevation threshold (see Methods). This modification enables the model to capture both the deterministic dispersal limits imposed by high-relief regions, which are shared by many species, and the stochastic nature of barrier formation due to species-specific constraints, providing a closer analogue to real biogeographic processes. Incorporating elevational constraints generates more realistic geographic patterns of range size on some continents (Fig. 4A–C), particularly in South America where the Andes represent a crucial barrier along which small ranges are concentrated, and partially also in Eurasia and Africa, although here the empirical spatial distribution of range sizes seems to additionally reflect the role of biome boundaries not included in the model. However, topography did not improve model performance in some groups, particularly in birds, likely because strong dispersal abilities make birds less constrained by elevation than other vertebrate groups. Furthermore, elevation-related model did not yield more realistic predictions for geographic range size patterns in Australia that lacks dominant mountain ridges, nor in North America, except for mammals (Table 1). Mountain ridges increase the local density of dispersal barriers, generating smaller ranges (similarly as the model with higher inner-barrier density), but it did not lead to a significantly improved fit of the frequency distribution of range sizes (*SI Appendix,* Fig. S2 and *SI Appendix*, Table S2).

**Fig. 4:**
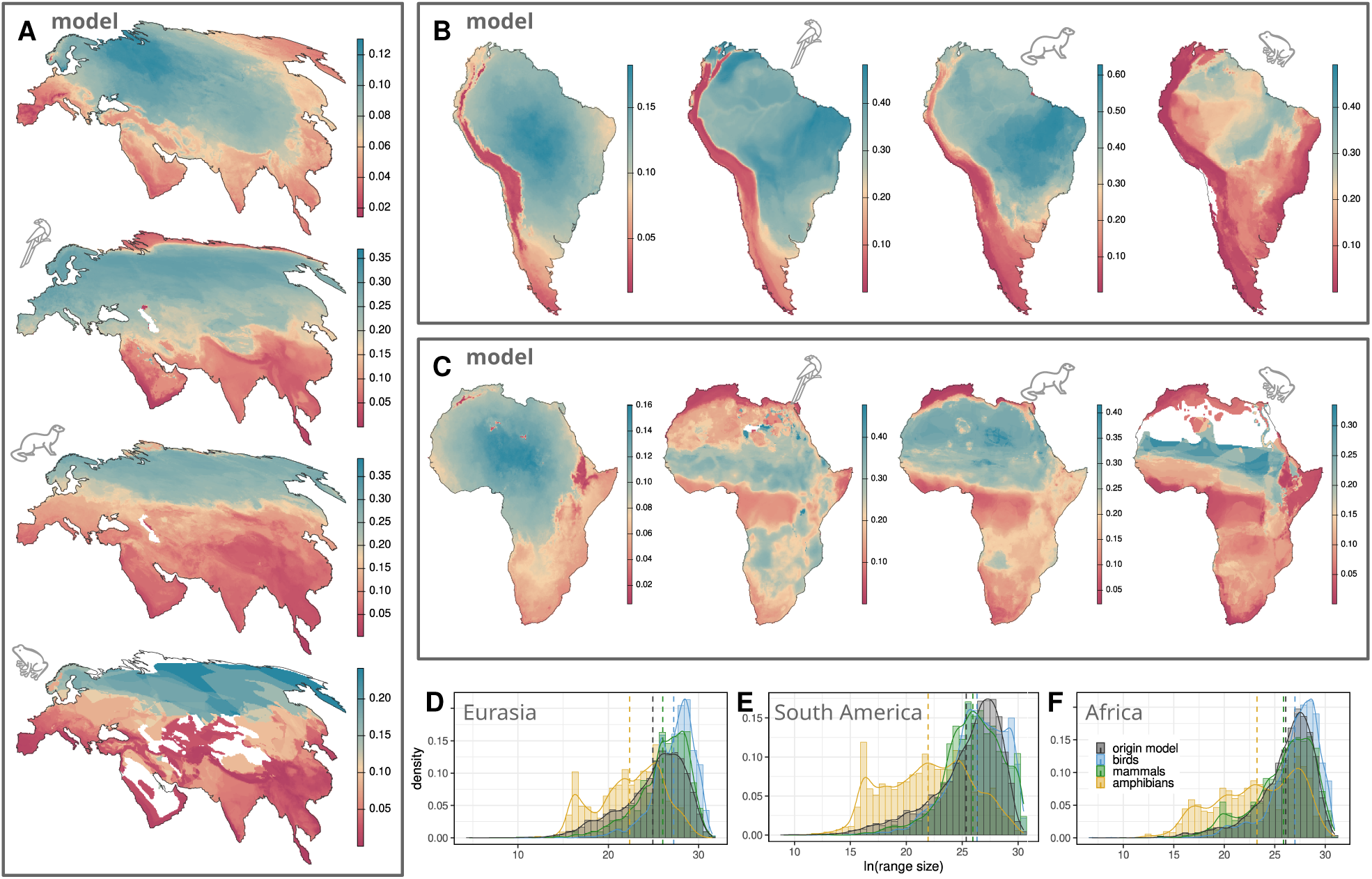
Adding realism to the models. (A, B, C) Geographic distribution of median range sizes predicted by the geometric model that includes elevation (the topographic model) compared with observed patterns for Eurasia (A), South America (B), and Africa (C) (red indicates smaller ranges, blue larger ranges, the color scale is adjusted to each continent separately). Model results are based on simulations of 5,000 species, rasterized at 30-km resolution in the Mollweide projection. Inner barriers are placed randomly with density Pois(*D* = 15), with additional barriers defined by elevational barriers. (D, E, F) Effect of productivity-dependent species origination on frequency distributions of range sizes. Frequency distributions of modeled (grey) and observed range sizes for birds (blue), mammals (green), and amphibians (yellow) in Eurasia (D), South America (E), and Africa (F) are shown on a logarithmic scale (m^2^). The inner-barrier density is Pois(*D* = 10). Altered species origination probability leads to a significantly improved fit to observed distributions. Solid lines represent kernel densities, and dashed lines indicate mean values.

### Productivity-dependent origin model

Variation in model fit among continents and taxa may partly reflect the model assumption that species can spread randomly from any point within the domain, ignoring unequal distribution of species richness across Earth’s surface, e.g. the latitudinal diversity gradient (LDG). This may explain why empirical geographic range patterns in North America reveal a strong south-north trend and relatively poorer model fit. However, when we modified the model so that the probability of the initiation of the species spreading was proportional to primary productivity, the major correlate of species richness (15, 16), predicted geographic patterns of range size changed only slightly (*SI Appendix*, Fig. S3). This indicates that the range size patterns are not a mere product of species richness gradient and instead emerge as a geometric consequence of the location of the barriers, the first-order patterns – namely the coincidence of species rarity with domain boundaries – being quite robust across alternative assumptions about species origination. However, this altered species origination significantly improved the fit to the observed frequency distributions of range sizes (Fig. 4D–F and *SI Appendix*, Fig. S4 and *SI Appendix*, Table S3). This improvement arose because centres of continents are typically less productive, and increasing probability of species origination near marginal regions elevates the proportion of small-ranged species. The advanced model thus showed that the exact shape of the frequency distribution of range sizes depends on the spatial distribution of species origination (see also ref. 17).

## Discussion

Our framework may appear similar to other geometric models that have been developed to explain diversity gradients via the mid-domain effect (MDE)—the higher overlap of species ranges in the middle of the domain (17–20). However, although our basic model that assumes random barrier location and equal probability of species origination across the whole area also produces highest species richness in the center of the domain (*SI Appendix*, Fig. S5 A and C), the overall logic is different. The MDE models assume a theoretical or empirical frequency distribution of range sizes and random reshuffling of whole ranges within a domain (15, 21–23). The mid-domain peak of species richness results from a necessary accumulation of wide-ranged species in the middle of the domain, and the spatial trends in range size depend on the assumed frequency distribution of range sizes (17, 19, 24, 25). Other geometric models, which allow ranges to extend across domain boundaries, can also reproduce broad spatial patterns in average range size, but are still based on pre-set range sizes (26–29). In contrast, our model assumes the opposite – range sizes emerge as an outcome of randomly positioned boundaries within the domain. The MDE framework has been criticized for its assumption that range sizes pre-exist, even though the environmental gradients that would necessarily shape them are precisely what the null model seeks to exclude (30–32). Because MDE models rely on a size frequency distribution which itself emerges from particular evolutionary or ecological processes, the model that pretends to be purely geometric in fact includes these processes. We argue that our model in which dispersal barriers are primary determinants of range size patterns represents a more parsimonious approach that can serve as a first approximation of geographic variability in range size. Interestingly, although the model that assumes a non-random productivity-dependent species origin does not typically improve the predictions of spatial range size patterns (*SI Appendix*, Fig. S3), it leads to very realistic size frequency distributions (Fig. 4D–F and *SI Appendix*, Fig. S4). Not surprisingly, it also leads to realistic species richness patterns (*SI Appendix*, Fig. S5 B and D), similarly as the MDE reproduces species richness patterns if it is complemented with non-random productivity-based spreading of the ranges within the domain (15).

Although our geometric model well predicts first-order macroecological patterns comprising range sizes, there are biological effects that necessarily play a role. First and most importantly, the model works only for particular densities of inner dispersal barriers, and these densities probably reflect biologically relevant (and perhaps taxon-specific) trade-offs. Specifically, too high density of barriers would imply a strong dependence of species distribution on environmental gradients, i.e. very low species environmental tolerance and high specialization. This would lead to a consequently high extinction risk, so that the system would not be sustainable in the long term. On the other hand, too low barrier density would appear only if most species were generalists that would be able to overcome spatially changing conditions. Such a system could not allow many species to coexist and could be broken by the speciation process. Second, the deviations from the model predictions in particular continents indicate that the location of barriers is often far from random, and the variability among taxa suggests that the role of specific biological traits may be essential even though the major barriers like coastlines still represent a key determinant of their range size.

## Conclusions

Species ranges and their limits result from a complex dynamic of species and their environment, and reflect phylogenetic history of respective lineages, including idiosyncratic sequences of speciation and extinction. Therefore, range size is certainly not a random species trait passively reflecting the distribution of dispersal barriers. However, our model indicates that the very existence of the geographic template with some barriers imposes an important constraint on macroecological range size patterns and their first-order form. Whenever species occur close to some of the hard, outer barriers, i.e. close to coastlines, on peninsulas or smaller land masses including islands, this very presence of the barrier means that their spread is constrained in some directions, increasing the probability that other, species-specific barriers will constrain their further dispersal and thus that their distribution will remain limited. Species living close to hard geographic barriers are thus naturally more endangered due to their inability to shift their ranges during environmental changes, and measures to prevent their extinction due to these changes represent an urgent task.

## Methods

### Basic model

The basic model consists of hard domain boundaries and probabilistic inner barriers defined on a real spheroid representation of the Earth. Domains correspond to five major continental land masses (Eurasia, Africa, North America, South America, and Australia), delineated by spherical polygons. Inner barriers are represented by randomly placed great circles, i.e. the largest circles that can be drawn on a sphere with planes passing through its center, with their number drawn from a Poisson distribution with mean density *D* (Pois(*D*)). Each great circle is defined by a random point on the sphere and a random initial azimuth determining the orientation of the circle. The simulations proceed until the required number of circles intersecting a given continent is reached, partitioning the domain into polygons.

A species originates at a random location within a continent, and its range is defined by the polygon containing the point of origin. Species range size is calculated as the area of the corresponding polygon on the sphere. For each simulation, 5,000 species were generated, each with an independent realization of inner barriers drawn from Pois(*D*). Resulting ranges were rasterized onto a Mollweide projection at 30-km resolution to derive geographic patterns of range size. For each grid cell, we calculated the median range size of overlapping species and the proportion of species with range sizes below the continental median. Frequency distributions of range sizes were obtained from all simulated species within each continent, and analyses were repeated separately for each continent and for different *D* values.

### Topographic model

The extended model incorporates topography by combining randomly placed inner barriers (mean density *D*) with elevation-based constraints derived from real-world data. For each continent, we have chosen one or two elevations that separated elevation bands, and each species range was assumed to be confined to just one elevation band based on the point of origin. The elevation barriers may thus truncate the polygon that was initially defined by the random geographic barriers. If this truncation produced multiple polygons comprising given elevation band within the initial polygon, total range size was calculated as the combined area of these fragments.

Elevational barriers were defined separately for each continent to capture broad topographic variability and represent approximate major elevational zones. We assumed that the barriers may be partly species-specific, so that they were drawn from the Normal distribution with continent-specific means: Eurasia, 691 m above sea level (a.s.l.) [species-specific barriers were drawn from N(691, 300)]; Africa and North America, 1000 and 2000 m a.s.l. [N(1000, 200) and N(2000, 200)]; South America, 1200 and 4000 m a.s.l. [N(1200, 300) and N(4000, 300)]; and Australia, 500 and 1000 m a.s.l. [N(500, 100) and N(1000, 100)].

Elevation data were obtained using the *geodata* package in R (33), based on Shuttle Radar Topography Mission (SRTM) data at 5′ resolution from https://srtm.csi.cgiar.org/.

### Productivity-dependent origin model

This model alters the assumption of uniform probability of species origination by linking the probability of origin to primary productivity, a key correlate of species richness (16). We used the Normalized Difference Vegetation Index (NDVI) as a proxy for productivity (34). For each polygon defined by random barriers, we calculated its area (*A*) and mean NDVI value (*P*). The probability of selecting a polygon as a species range was proportional to log(*A*) × *P*, rather than being determined by a random point.

In the combined model incorporating both non-uniform origin and topography, polygons were first selected according to the NDVI-weighted probabilities, and then a random point within the selected polygon determined the applicable elevation band defined by elevational barriers, from which the range size was calculated. The results of the combined model are shown in Table 1. NDVI data were obtained from the MODIS MOD13A3.v061 vegetation index product (35), providing monthly NDVI values for the period of 2000-2024 at 1000-m resolution in a custom sinusoidal projection (https://modis.gsfc.nasa.gov/data/dataprod/mod13.php). Data were accessed via the AppEEARS platform (https://appeears.earthdatacloud.nasa.gov). We removed bad quality observations based on MODIS VI quality and reliability layers (see MODIS VI User’s Guide https://vip.arizona.edu/MODIS_UsersGuide.php and R scripts in *SI Software*). The overall mean NDVI was calculated by averaging the 25-year means of the 12 calendar months. The final raster was reprojected to WGS84 at 0.1° resolution for each continent.

### Species ranges

Spatial range data for mammals and amphibians were obtained from the IUCN Red List of Threatened Species (36) (www.iucnredlist.org), and for birds from BirdLife International (37) https://datazone.birdlife.org). Species distributions were based on extent-of-occurrence (EOO) polygon coding following IUCN mapping standards (see the IUCN Red List Categories and Criteria www.iucnredlist.org/resources/mappingstandards and R scripts in *SI Software*). We included only breeding bird ranges and excluded purely marine species. Analyses were restricted to ranges overlapping continental mainlands.

Empirical range sizes were calculated from spatial polygons which were subsequently rasterized onto a Mollweide projection at 30-km resolution to derive geographic patterns of range size. For each grid cell, we calculated the median range size of overlapping species and the proportion of species with range sizes below the continental first three quartiles. Frequency distributions of range sizes were obtained from all species within each taxon within each continent.

We quantified the correspondence between observed and model-predicted patterns using ordinary least squares (OLS) linear regression. For each continent and taxon, the observed median range size per grid cell was treated as the dependent variable and the model-predicted median range size as the explanatory variable. Model performance was evaluated using the adjusted coefficient of determination (R²), which accounts for differences in sample size and provides a measure of explained variance. Analyses were conducted separately for each model variant, continent and taxon. The results are shown in Table 1.

All analyses were performed in R version 4.3.0 (2023-04-21) (R Core Team) (33).

## Data and Software Availability

Species distribution data for mammals and amphibians are available from the IUCN Red List of Threatened Species (https://www.iucnredlist.org). Bird distribution data are available from BirdLife International (https://datazone.birdlife.org). Shuttle Radar Topography Mission (SRTM) elevation data are accessible via CGIAR-CSI (https://srtm.csi.cgiar.org) and the *geodata* R package. NASA MODIS MOD13A3 vegetation index (NDVI) data are available at https://modis.gsfc.nasa.gov/data/dataprod/mod13.php.

All custom R scripts required for data processing, simulations, statistical analyses and figures are provided in *SI Software*.

## Author Contributions

A.T. and D.S. conceptualized the study, conducted the investigation, and wrote the manuscript. A.T. developed the methodology, performed all data analyses, and created the visualizations.

## Acknowledgments

This study was supported by the Czech Science Foundation grant GA ČR 25-18055S.

We further thank Petr Keil for his help with conceptualization of the core idea of the study and Roman Kotecký for discussions. We also thank our colleagues who encouraged us in this work with their feedback.

---

**Fig. S1:**
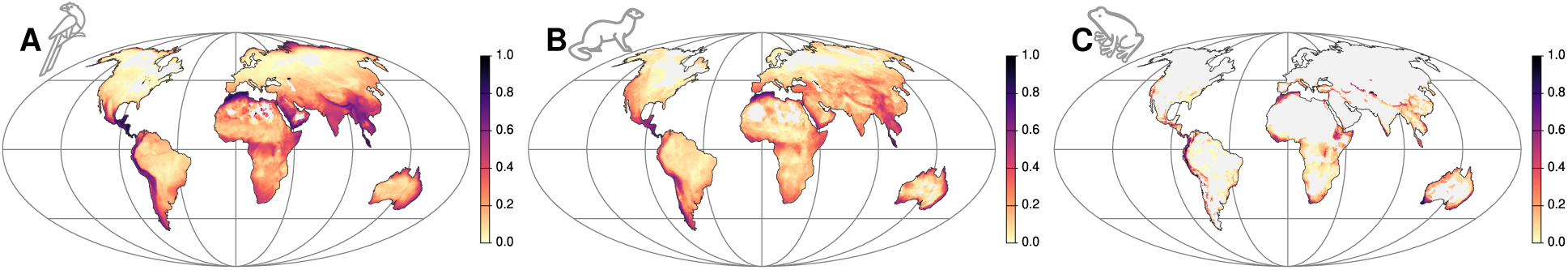
Geographic patterns of small-ranged vertebrate species. Geographic patterns of the proportion of small-ranged species (first three quartiles), for birds (A), mammals (B), and amphibians (C). Maps are displayed in the Mollweide projection and rasterized at 30-km resolution.

**Fig. S2:**
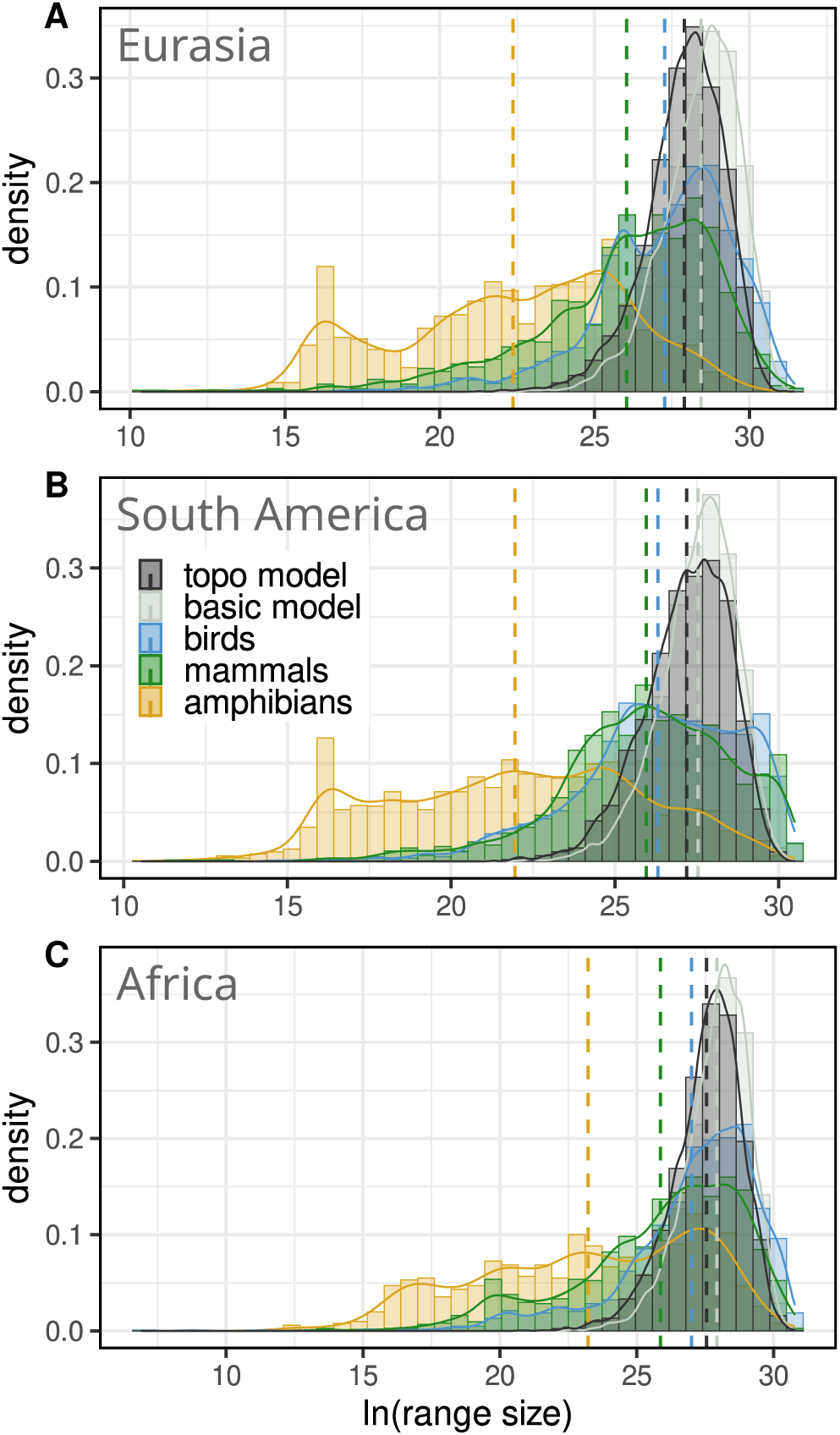
Effect of topography on frequency distributions of range sizes. Frequency distributions of modeled (grey) and observed range sizes for birds (blue), mammals (green), and amphibians (yellow) in Eurasia (A), South America (B), and Africa (C), shown on a logarithmic scale (m^2^). Two variants of the model are shown: the basic model (light grey) with inner-barrier density Pois(*D* = 15) and the topographic model (dark grey) with inner-barrier density Pois(*D* = 15) with additional elevational barriers. Incorporating topography has only a small effect on the size frequency distribution in comparison to the basic model. Solid lines represent kernel densities, and dashed lines indicate mean values.

**Fig. S3:**
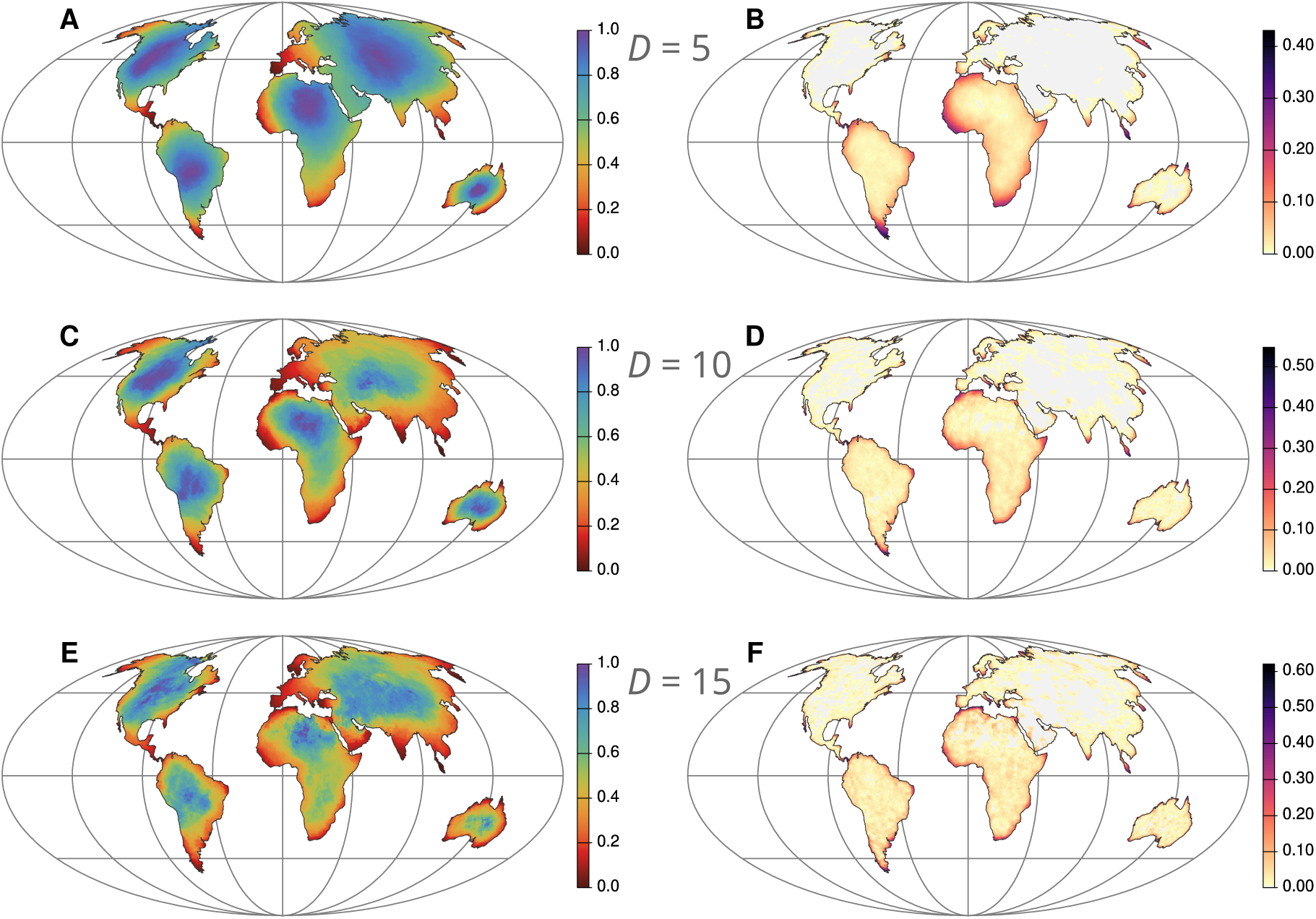
Effect of productivity-dependent species origination on geographic patterns of range size. (A, C, E) Geographic patterns of median range size, scaled separately within each continent (red indicates smaller ranges, blue larger ranges). (B, D, F) Geographic patterns of the proportion of small-ranged species (below the continental median). (A to F) Results are based on simulations of 5,000 species and rasterized to 30-km grid cells in the Mollweide projection. Inner barriers are placed randomly with densities Pois(*D* = 5) (A and B), Pois(*D* = 10) (C and D), and Pois(*D* = 15) (E and F). The probability of species origination is proportional to primary productivity. Despite this modification, the spatial patterns remain similar to those of the basic model.

**Fig. S4:**
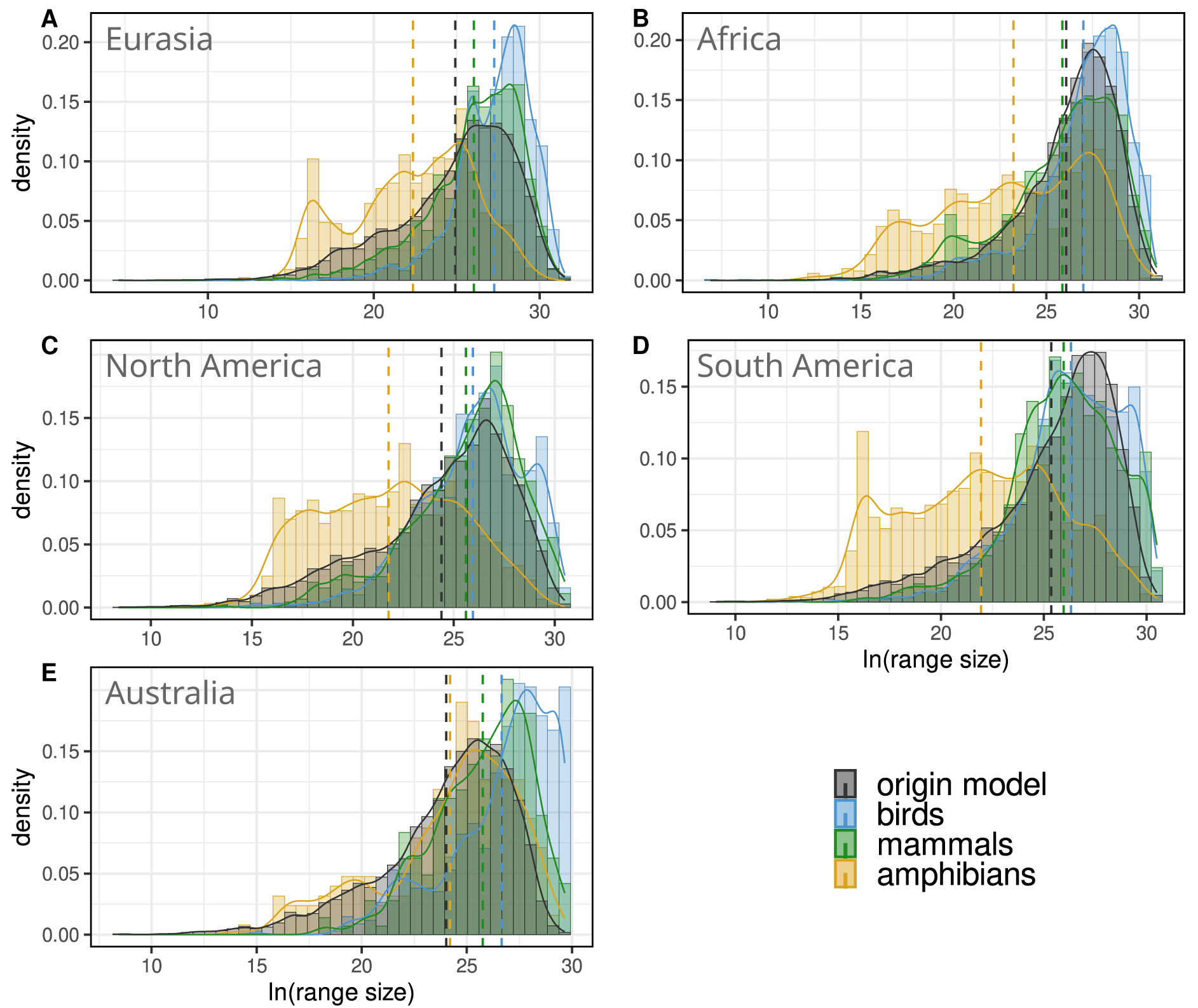
Effect of productivity-dependent species origination on frequency distributions of range sizes. Frequency distributions of modeled (grey) and observed range sizes for birds (blue), mammals (green), and amphibians (yellow) in Eurasia (A), Africa (B), North America (C), South America (D), and Australia (E), shown on a logarithmic scale (m^2^). The productivity-dependent origin model (grey) has inner-barrier density Pois(*D* = 10). Altered species origination leads to a significantly improved fit to observed distributions. Solid lines represent kernel densities, and dashed lines indicate mean values.

**Fig. S5:**
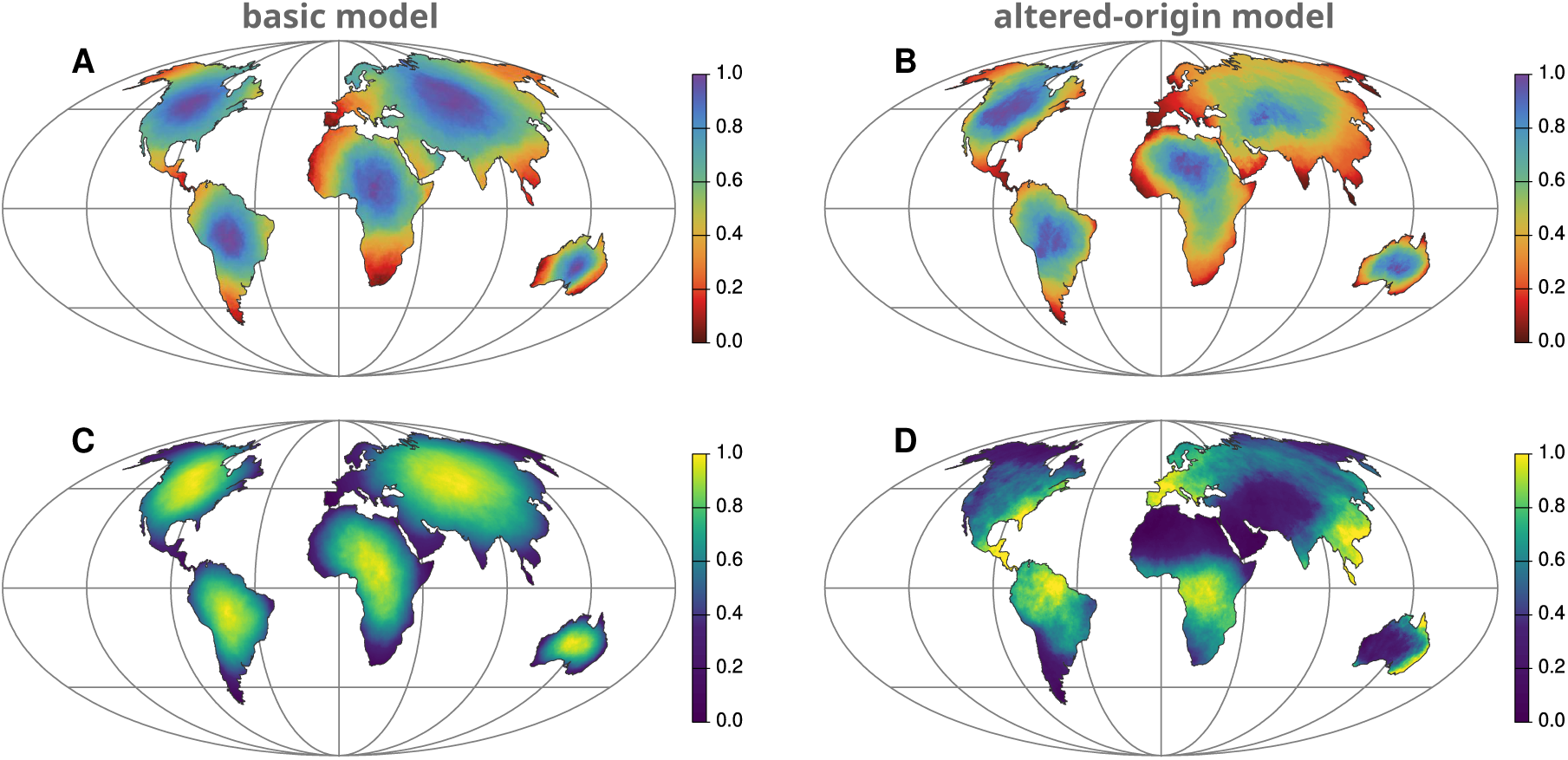
Geographic patterns of species richness of the basic and the productivity-dependent species origination model. (A and B) Geographic patterns of median range size of the basic model (A) and the productivity-dependent species origination model (B), scaled separately within each continent (red indicates smaller ranges, blue larger ranges). (C and D) Geographic patterns of species richness of the basic model (C) and the productivity-dependent species origination model (D). Altered species origination changes spatial patterns of range size only slightly. The results of both models are based on simulations of 5,000 species and rasterized to 30-km grid cells in the Mollweide projection. Inner barriers were placed randomly with densities Pois(*D* = 10).

**Table S1:**
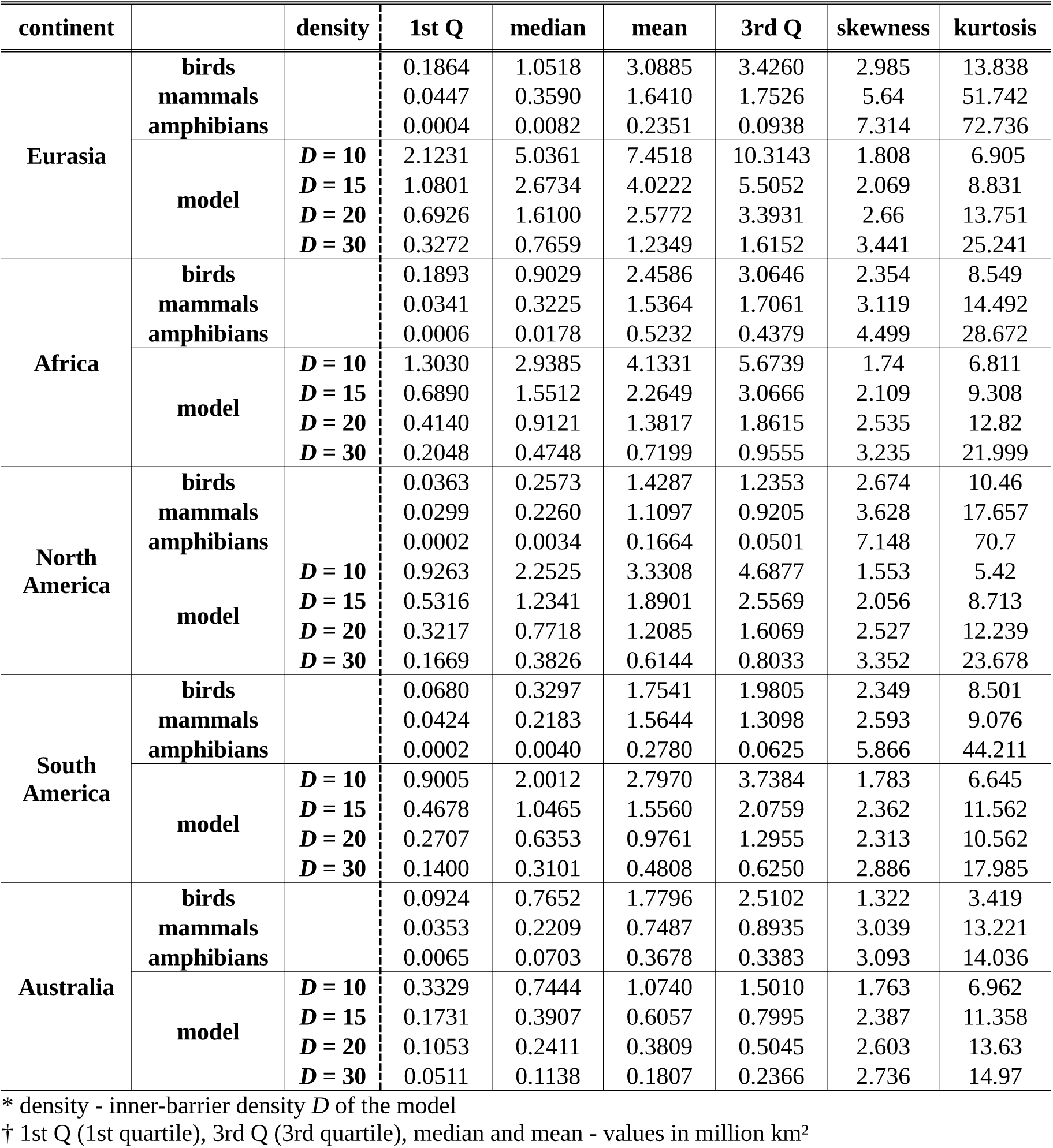
Frequency distributions of range sizes of the basic model and observed in vertebrate taxa. Statistics describing the shape of size frequency distributions of modeled and observed (in birds, mammals, and amphibians) ranges across individual continents.

| continent |  | density | 1st Q | median | mean | 3rd Q | skewness | kurtosis |
| --- | --- | --- | --- | --- | --- | --- | --- | --- |
| Eurasia | birds |  | 0.1864 | 1.0518 | 3.0885 | 3.4260 | 2.985 | 13.838 |
|  | mammals |  | 0.0447 | 0.3590 | 1.6410 | 1.7526 | 5.64 | 51.742 |
|  | amphibians |  | 0.0004 | 0.0082 | 0.2351 | 0.0938 | 7.314 | 72.736 |
| | model | $D = 10$ | 2.1231 | 5.0361 | 7.4518 | 10.3143 | 1.808 | 6.905 |
| | | $D = 15$ | 1.0801 | 2.6734 | 4.0222 | 5.5052 | 2.069 | 8.831 |
| | | $D = 20$ | 0.6926 | 1.6100 | 2.5772 | 3.3931 | 2.66 | 13.751 |
| | | $D = 30$ | 0.3272 | 0.7659 | 1.2349 | 1.6152 | 3.441 | 25.241 |
| Africa | birds |  | 0.1893 | 0.9029 | 2.4586 | 3.0646 | 2.354 | 8.549 |
|  | mammals |  | 0.0341 | 0.3225 | 1.5364 | 1.7061 | 3.119 | 14.492 |
|  | amphibians |  | 0.0006 | 0.0178 | 0.5232 | 0.4379 | 4.499 | 28.672 |
| | model | $D = 10$ | 1.3030 | 2.9385 | 4.1331 | 5.6739 | 1.74 | 6.811 |
| | | $D = 15$ | 0.6890 | 1.5512 | 2.2649 | 3.0666 | 2.109 | 9.308 |
| | | $D = 20$ | 0.4140 | 0.9121 | 1.3817 | 1.8615 | 2.535 | 12.82 |
| | | $D = 30$ | 0.2048 | 0.4748 | 0.7199 | 0.9555 | 3.235 | 21.999 |
| North America | birds |  | 0.0363 | 0.2573 | 1.4287 | 1.2353 | 2.674 | 10.46 |
|  | mammals |  | 0.0299 | 0.2260 | 1.1097 | 0.9205 | 3.628 | 17.657 |
|  | amphibians |  | 0.0002 | 0.0034 | 0.1664 | 0.0501 | 7.148 | 70.7 |
| | model | $D = 10$ | 0.9263 | 2.2525 | 3.3308 | 4.6877 | 1.553 | 5.42 |
| | | $D = 15$ | 0.5316 | 1.2341 | 1.8901 | 2.5569 | 2.056 | 8.713 |
| | | $D = 20$ | 0.3217 | 0.7718 | 1.2085 | 1.6069 | 2.527 | 12.239 |
| | | $D = 30$ | 0.1669 | 0.3826 | 0.6144 | 0.8033 | 3.352 | 23.678 |
| South America | birds |  | 0.0680 | 0.3297 | 1.7541 | 1.9805 | 2.349 | 8.501 |
|  | mammals |  | 0.0424 | 0.2183 | 1.5644 | 1.3098 | 2.593 | 9.076 |
|  | amphibians |  | 0.0002 | 0.0040 | 0.2780 | 0.0625 | 5.866 | 44.211 |
| | model | $D = 10$ | 0.9005 | 2.0012 | 2.7970 | 3.7384 | 1.783 | 6.645 |
| | | $D = 15$ | 0.4678 | 1.0465 | 1.5560 | 2.0759 | 2.362 | 11.562 |
| | | $D = 20$ | 0.2707 | 0.6353 | 0.9761 | 1.2955 | 2.313 | 10.562 |
| | | $D = 30$ | 0.1400 | 0.3101 | 0.4808 | 0.6250 | 2.886 | 17.985 |
| Australia | birds |  | 0.0924 | 0.7652 | 1.7796 | 2.5102 | 1.322 | 3.419 |
|  | mammals |  | 0.0353 | 0.2209 | 0.7487 | 0.8935 | 3.039 | 13.221 |
|  | amphibians |  | 0.0065 | 0.0703 | 0.3678 | 0.3383 | 3.093 | 14.036 |
| | model | $D = 10$ | 0.3329 | 0.7444 | 1.0740 | 1.5010 | 1.763 | 6.962 |
| | | $D = 15$ | 0.1731 | 0.3907 | 0.6057 | 0.7995 | 2.387 | 11.358 |
| | | $D = 20$ | 0.1053 | 0.2411 | 0.3809 | 0.5045 | 2.603 | 13.63 |
| | | $D = 30$ | 0.0511 | 0.1138 | 0.1807 | 0.2366 | 2.736 | 14.97 |

\* density - inner-barrier density $D$ of the model
† 1st Q (1st quartile), 3rd Q (3rd quartile), median and mean - values in million km<sup>2</sup>

**Table S2:** Frequency distributions of range sizes of the topographic model. Statistics describing the shape of size frequency distributions of ranges predicted by the topographic model under varying inner-barrier densities across individual continents.

| continent | density | 1st Q | median | mean | 3rd Q | skewness | kurtosis |
| --- | --- | --- | --- | --- | --- | --- | --- |
| Eurasia | $D = 10$ | 1.0770 | 2.7276 | 4.3876 | 5.9620 | 2.091 | 8.582 |
| | $D = 15$ | 0.6413 | 1.4861 | 2.4003 | 3.19 | 2.492 | 13.214 |
| | $D = 20$ | 0.399 | 0.9561 | 1.5745 | 2.0435 | 3.072 | 20.601 |
| Africa | $D = 10$ | 0.7835 | 1.8923 | 2.9864 | 4.0998 | 2.105 | 9.014 |
| | $D = 15$ | 0.4519 | 1.0766 | 1.7149 | 2.2436 | 2.68 | 13.928 |
| | $D = 20$ | 0.2665 | 0.6598 | 1.0743 | 1.4013 | 2.433 | 11.29 |
| North America | $D = 10$ | 0.5311 | 1.2298 | 2.1133 | 2.8208 | 2.056 | 7.857 |
| | $D = 15$ | 0.2946 | 0.7188 | 1.2199 | 1.5675 | 2.664 | 13.082 |
| | $D = 20$ | 0.2108 | 0.4968 | 0.819 | 1.0623 | 2.827 | 15.116 |
| South America | $D = 10$ | 0.5481 | 1.4763 | 2.2547 | 3.1158 | 1.808 | 6.915 |
| | $D = 15$ | 0.2912 | 0.7414 | 1.2503 | 1.7062 | 2.371 | 10.861 |
| | $D = 20$ | 0.1975 | 0.4824 | 0.8156 | 1.0731 | 2.752 | 14.873 |
| Australia | $D = 10$ | 0.2309 | 0.5612 | 0.8738 | 1.2066 | 2.051 | 8.462 |
| | $D = 15$ | 0.1255 | 0.3076 | 0.487 | 0.6692 | 2.14 | 9.328 |
| | $D = 20$ | 0.0828 | 0.1953 | 0.308 | 0.4108 | 2.372 | 11.049 |

\* density - inner-barrier density $D$ of the model
† 1st Q (1st quartile), 3rd Q (3rd quartile), median and mean - values in million km<sup>2</sup>

**Table S3:** Frequency distributions of range sizes of the altered-origin model. Statistics describing the shape of size frequency distributions of ranges predicted by the productivity-dependent origin model under varying inner-barrier densities across individual continents.

| continent | density | 1st Q | median | mean | 3rd Q | skewness | kurtosis |
| --- | --- | --- | --- | --- | --- | --- | --- |
| Eurasia | $D = 10$ | 0.0092 | 0.1446 | 1.2726 | 0.9983 | 5.479 | 46.491 |
| | $D = 15$ | 0.0104 | 0.1221 | 0.765 | 0.6429 | 5.633 | 50.624 |
| | $D = 20$ | 0.0103 | 0.0916 | 0.4753 | 0.4371 | 5.228 | 43.819 |
| Africa | $D = 10$ | 0.059 | 0.4105 | 1.4208 | 1.5439 | 3.853 | 23.159 |
| | $D = 15$ | 0.0323 | 0.2029 | 0.6975 | 0.7633 | 4.058 | 26.461 |
| | $D = 20$ | 0.022 | 0.1204 | 0.3849 | 0.4262 | 4.306 | 32.177 |
| North America | $D = 10$ | 0.0057 | 0.085 | 0.6507 | 0.531 | 4.576 | 30.753 |
| | $D = 15$ | 0.005 | 0.0616 | 0.3609 | 0.3248 | 5.494 | 48.75 |
| | $D = 20$ | 0.0054 | 0.0451 | 0.2368 | 0.2177 | 7.633 | 105.629 |
| South America | $D = 10$ | 0.0227 | 0.2381 | 0.9353 | 1.0293 | 3.714 | 21.592 |
| | $D = 15$ | 0.0165 | 0.1243 | 0.458 | 0.4809 | 4.493 | 32.665 |
| | $D = 20$ | 0.0106 | 0.0676 | 0.2635 | 0.272 | 4.156 | 27.516 |
| Australia | $D = 10$ | 0.0059 | 0.0533 | 0.2654 | 0.2672 | 4.811 | 39.101 |
| | $D = 15$ | 0.0037 | 0.032 | 0.1385 | 0.1354 | 5.335 | 47.507 |
| | $D = 20$ | 0.0027 | 0.021 | 0.0816 | 0.0827 | 4.651 | 32.815 |

\* density - inner-barrier density $D$ of the model
† 1st Q (1st quartile), 3rd Q (3rd quartile), median and mean - values in million km<sup>2</sup>

## References

1. K. J. Gaston, The Structure and Dynamics of Geographic Ranges (Oxford University Press, 2003).

2. J. Smyèka, A. Toszogyova, D. Storch, The relationship between geographic range size and rates of species diversification. Nat Commun 14, 5559 (2023).

3. A. Alzate et al., Evolutionary age correlates with range size across plants and animals. Nat Commun 16, 7894 (2025).

4. J. H. Brown, G. C. Stevens, D. M. Kaufman, The Geographic Range: Size, Shape, Boundaries, and Internal Structure. Annual Review of Ecology and Systematics 27, 597–623 (1996).

5. K. J. Gaston, Species-range size distributions: products of speciation, extinction and transformation. Philos Trans R Soc Lond B Biol Sci 353, 219–230 (1998).

6. K. J. Gaston, T. Blackburn, Pattern And Process In Macroecology (Blackwell Publishing Ltd, 2000).

7. K. J. Gaston, R. G. Davies, C. E. Gascoigne, M. Williamson, The structure of global species–range size distributions: raptors & owls. Global Ecology and Biogeography 14, 67–76 (2005).

8. K. J. Gaston, Species-range-size distributions: patterns, mechanisms and implications. Trends in Ecology & Evolution 11, 197–201 (1996).

9. H. T. Arita, Range size in mid-domain models of species diversity. Journal of Theoretical Biology 232, 119–126 (2005).

10. G. C. Stevens, The Latitudinal Gradient in Geographical Range: How so Many Species Coexist in the Tropics. The American Naturalist 133, 240–256 (1989).

11. C. D. L. Orme et al., Global Patterns of Geographic Range Size in Birds. PLOS Biology 4, e208 (2006).

12. T. J. Davies, A. Purvis, J. L. Gittleman, Quaternary Climate Change and the Geographic Ranges of Mammals. The American Naturalist 174, 297–307 (2009).

13. G. Murali, R. Gumbs, S. Meiri, U. Roll, Global determinants and conservation of evolutionary and geographic rarity in land vertebrates. Sci Adv 7, eabe5582 (2021).

14. P. Keil, D. Storch, W. Jetz, On the decline of biodiversity due to area loss. Nat Commun 6, 8837 (2015).

15. D. Storch et al., Energy, range dynamics and global species richness patterns: reconciling mid-domain effects and environmental determinants of avian diversity. Ecology Letters 9, 1308–1320 (2006).

16. E. Bohdalková, A. Toszogyova, I. Šímová, D. Storch, Universality in biodiversity patterns: variation in species–temperature and species–productivity relationships reveals a prominent role of productivity in diversity gradients. Ecography 44, 1366–1378 (2021).

17. R. K. Colwell, G. C. Hurtt, Nonbiological Gradients in Species Richness and a Spurious Rapoport Effect. The American Naturalist 144, 570–595 (1994).

18. D. C. Lees, C. Kremen, L. Andriamampianina, A null model for species richness gradients: bounded range overlap of butterflies and other rainforest endemics in Madagascar. Biol J Linn Soc 67, 529–584 (1999).

19. R. K. Colwell, D. C. Lees, The mid-domain effect: geometric constraints on the geography of species richness. Trends in Ecology & Evolution 15, 70–76 (2000).

20. J. A. Grytnes, Ecological interpretations of the mid-domain effect. Ecology Letters 6, 883–888 (2003).

21. W. Jetz, C. Rahbek, Geometric constraints explain much of the species richness pattern in African birds. Proceedings of the National Academy of Sciences 98, 5661–5666 (2001).

22. P. Koleff, K. J. Gaston, Latitudinal gradients in diversity: real patterns and random models. Ecography 24, 341–351 (2001).

23. C. M. McCain, North American Desert Rodents: A Test of the Mid-Domain Effect in Species Richness. J Mammal 84, 967–980 (2003).

24. R. K. Colwell, C. Rahbek, N. J. Gotelli, The Mid-Domain Effect and Species Richness Patterns: What Have We Learned So Far? The American Naturalist 163, E1–E23 (2004).

25. R. K. Colwell et al., Peaks, plateaus, canyons, and craters: the complex geometry of simple mid-domain effect models. Evol Ecol Res 11, 355–370 (2009).

26. J. A. Grytnes, O. R. Vetaas, Species Richness and Altitude: A Comparison between Null Models and Interpolated Plant Species Richness along the Himalayan Altitudinal Gradient, Nepal. The American Naturalist 159, 294–304 (2002).

27. B. S. Sandel, M. J. McKone, Reconsidering null models of diversity: Do geometric constraints on species ranges necessarily cause a mid-domain effect? Diversity and Distributions 12, 467–474 (2006).

28. J. A. Grytnes, J. H. Beaman, T. S. Romdal, C. Rahbek, The mid-domain effect matters: simulation analyses of range-size distribution data from Mount Kinabalu, Borneo. Journal of Biogeography 35, 2138–2147 (2008).

29. B. Sandel, Geometric constraint model selection – an example with New World birds and mammals. Ecography 32, 1001–1010 (2009).

30. H. Laurie, J. A. Silander, Geometric Constraints and Spatial Pattern of Species Richness: Critique of Range-Based Null Models. Diversity and Distributions 8, 351–364 (2002).

31. F. A. Zapata, K. J. Gaston, S. L. Chown, Mid-domain models of species richness gradients: assumptions, methods and evidence. Journal of Animal Ecology 72, 677–690 (2003).

32. B. A. Hawkins, J. A. F. Diniz-Filho, A. E. Weis, The mid-domain effect and diversity gradients: is there anything to learn? Am Nat 166, E140–143 (2005).

33. R Core Team. R: A language and environment for statistical computing https://www.R-project.org (R Foundation for Statistical Computing, 2023).

34. A. Toszogyova, D. Storch, Global diversity patterns are modulated by temporal fluctuations in primary productivity. Global Ecology and Biogeography 28, 1827–1838 (2019).

35. K. Didan, MODIS/Terra Vegetation Indices Monthly L3 Global 1km SIN Grid V061 (MOD13A3) data set. LP DAAC 10.5067/MODIS/MOD13A3.061 (2021).

36. IUCN. The IUCN Red List of Threatened Species, Version 2024-2 https://www.iucnredlist.org (2024).

37. BirdLife International and Handbook of the Birds of the World. Bird species distribution maps of the world, Version 2022.2 http://datazone.birdlife.org/species/requestdis (2022).

